# FLASH proton reirradiation, with or without hypofractionation, mitigates chronic toxicity in the normal murine intestine, skin, and bone

**DOI:** 10.1101/2024.07.08.602528

**Authors:** Ioannis I. Verginadis, Anastasia Velalopoulou, Michele M. Kim, Kyle Kim, Ioannis Paraskevaidis, Brett Bell, Seyyedeh Azar Oliaei Motlagh, Antoneta Karaj, Esha Banerjee, Giovanni Finesso, Charles-Antoine Assenmacher, Enrico Radaelli, Jiawei Lu, Yuewei Lin, Mary E. Putt, Eric S. Diffenderfer, Chandan Guha, Ling Qin, James M. Metz, Amit Maity, Keith A. Cengel, Constantinos Koumenis, Theresa M. Busch

## Abstract

**Background and purpose:** The normal tissue sparing afforded by FLASH radiotherapy (RT) is being intensely investigated for potential clinical translation. Here, we studied the effects of FLASH proton RT (F-PRT) in the reirradiation setting, with or without hypofractionation. Chronic toxicities in three murine models of normal tissue toxicity including the intestine, skin, and bone were investigated.

**Materials and methods:** In studies of the intestine, single-dose irradiation was performed with 12 Gy of Standard proton RT (S-PRT), followed by a second dose of 12 Gy of F-PRT or S-PRT. Additionally, a hypofractionation scheme was applied in the reirradiation setting (3 x 6.4 Gy of F-PRT or S-PRT, given every 48 hrs). In studies of skin/bone of the murine leg, 15 Gy of S-PRT was followed by hypofractionated reirradiation with F-PRT or S-PRT (3 x 11 Gy).

**Results:** Compared to reirradiation with S-PRT, F-PRT reduced intestinal fibrosis and collagen deposition in the reirradiation setting and significantly increased survival rate, demonstrating its protective effects on intestinal tissues. In previously irradiated leg tissues, reirradiation with hypofractionated F-PRT created transient dermatitis that fully resolved in contrast to reirradiation with hypofractionated S-PRT. Lymphedema was also alleviated after a second course of radiation with F-PRT, along with significant reductions in the accumulation of fibrous connective tissue in the skin compared to mice reirradiated with S-PRT. The delivery of a second course of fractionated S-PRT induced tibial fractures in 83.3% of the mice, whereas only 20% of mice reirradiated with F-PRT presented with fractures.

**Conclusion:** These studies provide the first evidence of the sparing effects of F-PRT, in the setting of hypofractionated reirradiation. The results support FLASH as highly relevant to the reirradiation regimen where it exhibits significant potential to minimize chronic complications for patients undergoing RT.

## Introduction

Reirradiation has an increasing role in the ongoing care of cancer patients as they survive longer due to improvements in systemic therapies. Patients who develop local-regional relapses are unlikely to be cured with chemotherapy or immunotherapy alone^1, 2^. In some cases, surgical excision is a potential option for these patients, but in others, surgical options may be limited.

Reirradiation presents a major challenge to the radiation oncologist, as it is well documented that high cumulative doses are associated with early and late severe morbidities^1, 3-5^. Reirradiation could be considered in the case of a local recurrence or for a new primary cancer that arises in a previously irradiated field. In this case, the cumulative dose that is delivered to the region is important because the initial dose that was delivered will contribute to the probability of an adverse toxicity. Modern techniques such as intensity-modulated radiation therapy (IMRT) and stereotactic body radiotherapy (SBRT) can provide better conformality to the tumor volume than standard photon delivery and have been used to minimize the associated morbidities of reirradiation^6-14^. Proton radiotherapy (RT) is also used in reirradiation because of its physical characteristics, as protons deposit most of their energy within a narrow range (Bragg peak) with little exit dose. As a result, normal tissues may be exposed to less radiation, potentially enhancing the therapeutic ratio^1, 15-20^. Notwithstanding the benefits of these highly conformal modalities in the reirradiation setting, normal tissues will still likely receive high cumulative doses of radiation, leading to significant toxicity^1, 7, 8, 19^.

Ultra-high dose rate RT at mean dose rates of ≥40 Gy/s, known as FLASH RT, can spare a wide spectrum of normal tissues while being equi-effective to standard dose rate radiation in controlling tumor growth ^21^. Moreover, the conformal physical dosimetric advantages of protons may be enhanced by the temporal benefits of high dose rate delivery to reduce the morbidities of reirradiation when performed with FLASH proton RT (F-PRT) compared to Standard proton RT (S-PRT). We hypothesize that reirradiation with F-PRT will be safer and less toxic to normal tissues than reirradiation with S-PRT. We study reirradiation with F-PRT versus S-PRT after an initial delivery of S-PRT to murine intestinal tissues and to the skin/bone of the murine hind leg. Survival and pathological manifestations of tissue injury, inflammation, lymphedema, fibrosis, and bone damage, pertinent to gastrointestinal and leg reirradiation have been evaluated. We report the first evidence that FLASH proton RT can significantly spare normal tissues from the damage of reirradiation.

## 2. Materials and Methods

### 2.1. Murine studies

All animal experiments have been reviewed and approved by the Institutional Animal Care and Use Committee (IACUC) of the University of Pennsylvania and husbandry is provided by University Laboratory Animal Resources (ULAR) in AALAC-accredited facilities. Both male and female C57BL/6J mice, 10-13 weeks old (Jackson Labs, Bar Harbor, ME), were used in *in vivo* experimental procedures, as specified. Mice were checked daily and euthanized upon onset of severe morbidity including hunched posture, social withdrawal, relative immobility, or apparent weight loss >20%. Isoflurane in medical air was used to anesthetize mice for procedures, including RT. Details on the experimental design and numbers per condition are given in Supplementary Material.

### 2.2. Proton Beam Delivery

An IBA Proteus Plus C230 Cyclotron (Louvain-la-Neuve, Belgium) was used to deliver 230 MeV (∼32 g/cm^2^ range) proton beams through the fixed beam line in a dedicated research room^22^. A double scattered system was used to deliver a uniform field of 2 x 2 cm^2^, for both the intestinal and the leg radiation studies. Field uniformity was verified by EBT3 Gafchromic film prior to each experiment. Beam control systems have been described previously^22^. Details are given in Supplementary Material. In brief, as shown in Figures 1a and 2a, irradiation of the whole abdomen was performed in a single anesthetized mouse at a time, as previously described^22^. The F-PRT dose rate was 127.39 ± 5.95 Gy/sec and the S-PRT dose rate was 0.79 ± 0.04 Gy/sec. For the studies of the skin and tibia, radiation was delivered on the right hind leg of the mice as shown in Figure 3a and have been described elsewhere^23^. The F-PRT dose rate was 106.5 ± 3.8 Gy/sec and the S-PRT dose rate was 0.7 ± 0.07 Gy/sec. In all treatments, FLASH and Standard RT were delivered as shoot-through transmission beams.

**Figure 1.**
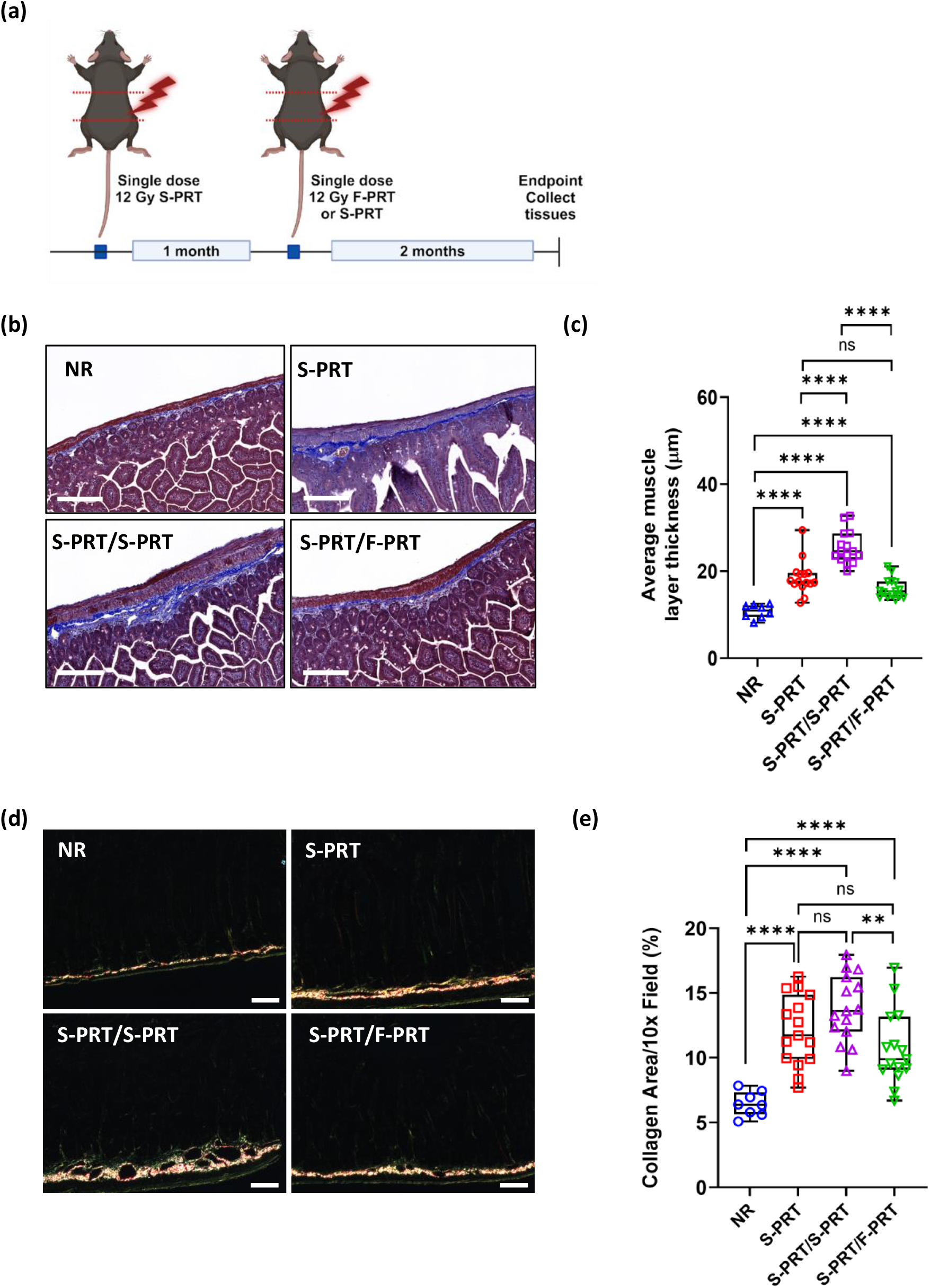
F-PRT spares intestinal fibrosis in a single-dose reirradiation preclinical model. **(a)** Schematic for single dose reirradiation study on the intestine. The mice received a single dose of 12 Gy of S-PRT on the entire abdomen, followed by another single dose of 12 Gy either of F-PRT or S-PRT, applied 1 month later. Red dotted lines indicate the irradiated area (Image was created with BioRender.com). **(b)** Representative images of Masson’s trichrome stain of intestinal sections, at 2 months after the second course of PRT; scale bar, 200 μm. **(c)** Box and whisker plot displays the muscle layer thickness (in μm). Each dot represents the average quantitative value per section per mouse (n = 8-15). **(d)** Representative images of Picrosirius red stain of intestinal sections, at 2 months after the second course of PRT; scale bar, 200 μm. **(e)** Box and whisker plot displays the % of fibrillar collagen per 10x field. Each dot represents the average quantitative value per section per mouse (n = 8-15). *P* values calculated with 2-tailed *t* test. **P<0.01, ****P<0.0001. Abbreviations: NR, no-RT (control); S-PRT, Standard proton RT; F-PRT, FLASH proton RT.

**Figure 2.**
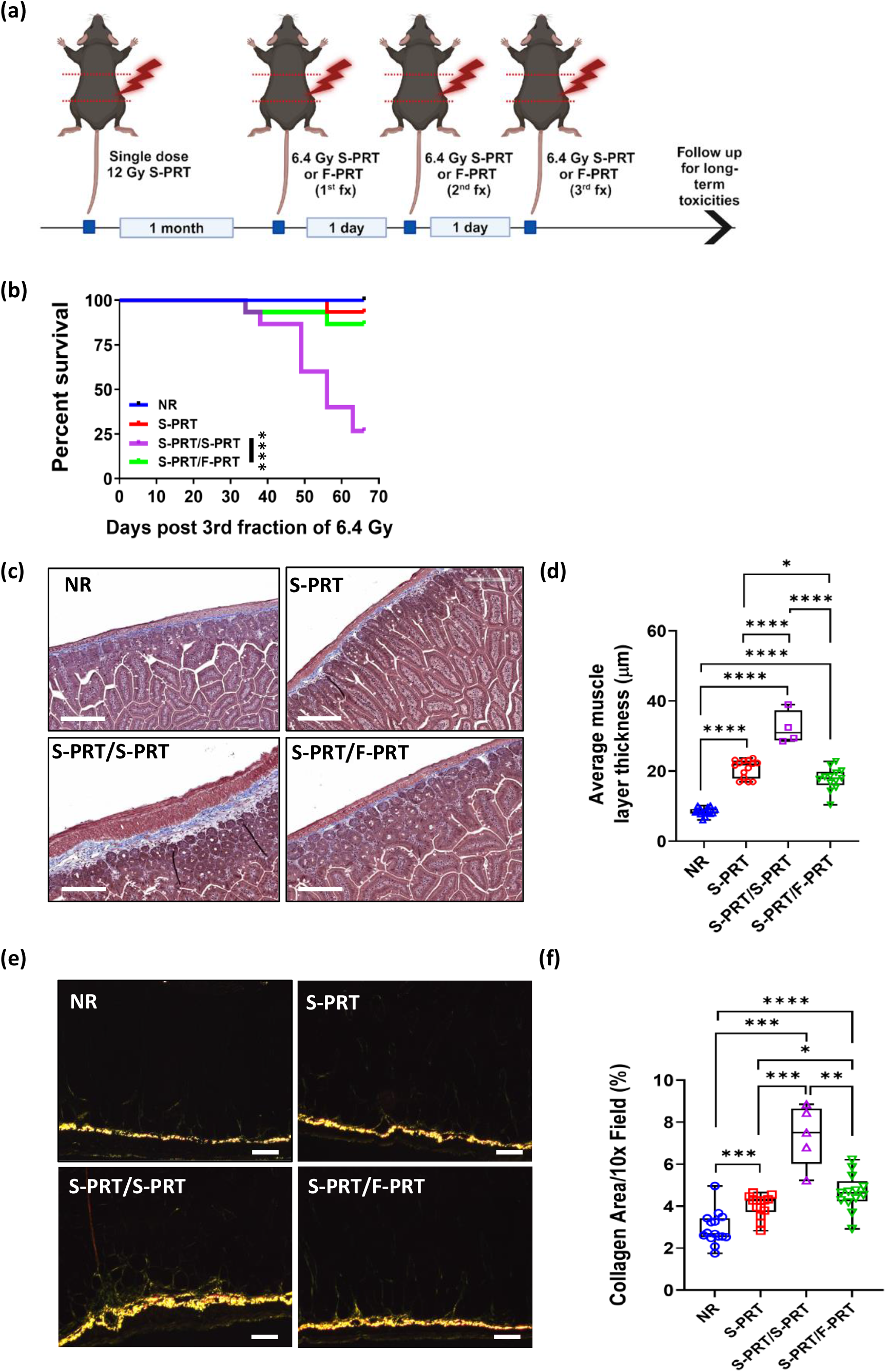
F-PRT provides a survival benefit and mitigates long-term intestinal toxicity following a hypofractionated reirradiation treatment scheme. **(a)** Schematic for fractionated reirradiation study on the intestine. The mice received a single dose of 12 Gy of S-PRT on the entire abdomen, followed by 3 fractions of 6.4 Gy of either F-PRT or S-PRT every other day, applied 1 month after the initial single dose irradiation. Red dotted lines indicate the irradiated area (Image was created with BioRender.com). **(b)** Kaplan–Meier survival analysis of the mice treated with hypofractionated reirradiation of the intestine. **(c)** Representative images of Masson’s trichrome stain of intestinal sections at 2 months post 3^rd^ fraction of 6.4 Gy of S-PRT or F-PRT; scale bar, 200 μm. **(d)** Box and whisker plot displays the muscle layer thickness (in μm). Each dot represents the average quantitative value per mouse (n = 4-15). **(e)** Representative images of picrosirius red stain of intestinal sections, at 2 months after the 3^rd^ fraction of 6.4 Gy of S-PRT or F-PRT; scale bar, 200 μm. **(f)** Box and whisker plot displays the % of fibrillar collagen per 10x field. Each dot represents the average quantitative value per section per mouse (n = 5-15). *P* values calculated with Log-rank (Mantel-Cox) test (for c and d) or 2-tailed *t* test (f. **P<0.01, ****P<0.0001. Abbreviations: NR, no-RT (control); S-PRT, Standard proton RT; F-PRT, FLASH proton RT.

**Figure 3.**
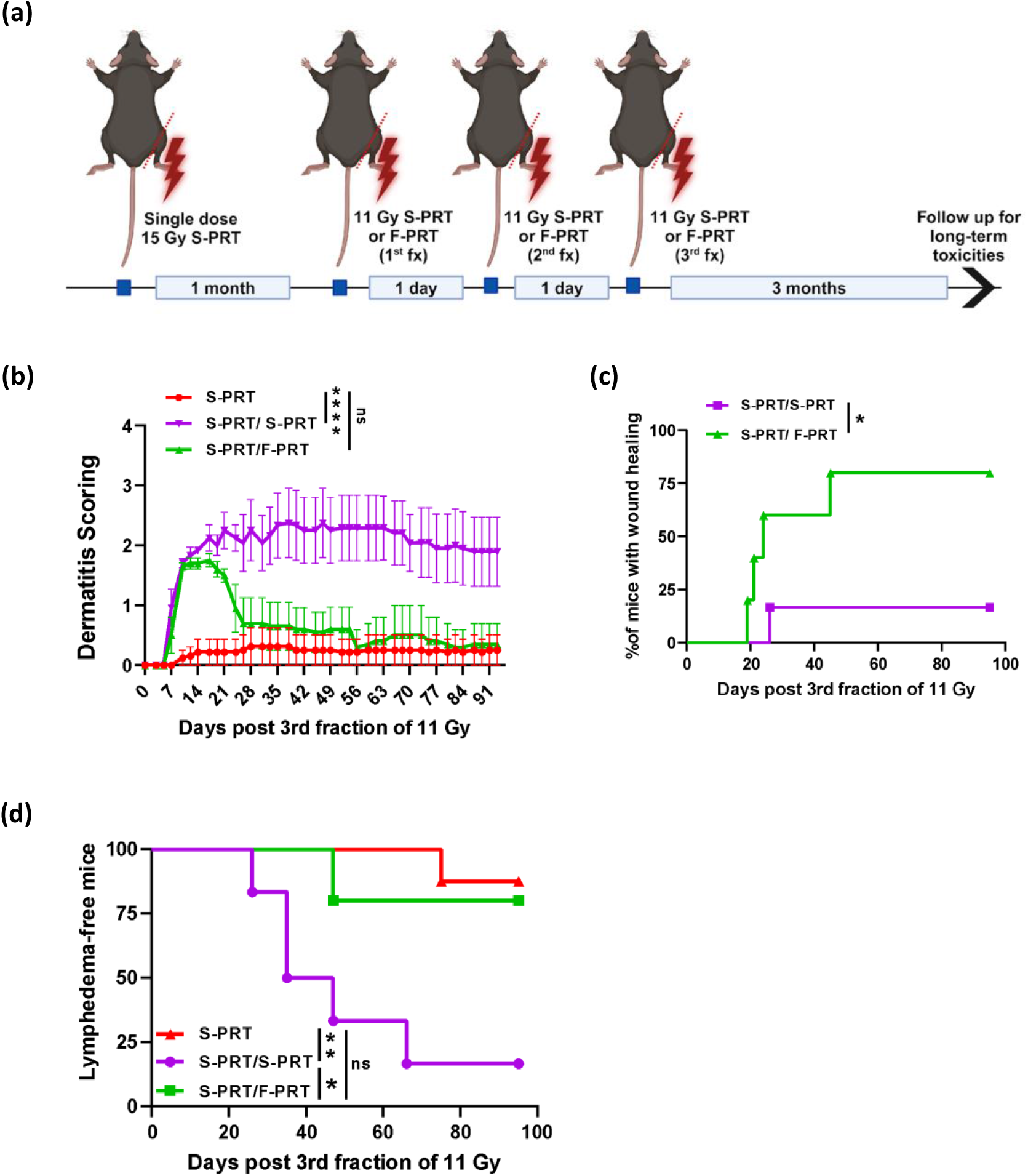
F-PRT mitigates skin dermatitis and lymphedema in a hypofractionated reirradiation treatment scheme. **(a)** Schematic for fractionated reirradiation study on the skin. The mice received a single dose of 15 Gy of S-PRT on the right hind leg, followed by 3 fractions of 11 Gy either of F-PRT or S-PRT, every other day, applied at 1 month after the initial single irradiation dose. Red dotted line indicates the irradiated area. **(b)** Line dot plot of dermatitis scoring across the treatment groups, throughout the monitoring period of 3 months after re-irradiation (observations are time matched for single irradiated animals). Data represent mean ± SEM (n=5-8) **(c)** Kaplan-Meier plot of the % of mice that experienced wound healing relative to time elapsed after the 3rd fraction of 11 Gy. **(d)** Kaplan-Meier plot of the % of lymphedema-free mice, relative to time elapsed after the 3rd fraction of 11 Gy. *P* values calculated with mixed-effect models, with a multiple comparison-Dunnett test (for b) or Log-rank (Mantel-Cox) test (for c and d). *P<0.05; **P<0.01; ****P<0.0001; ns, not significant. Abbreviations: NR, no-RT (control); S-PRT, Standard proton RT; F-PRT, FLASH proton RT.

### 2.3. Histopathology and quantification of fibrosis

Histopathology of skin and intestinal tissues was performed following standard protocols of the Comparative Pathology Core, at the School of Veterinary Medicine of University of Pennsylvania, for the purpose of quantifying fibrosis. Details are given in Supplementary Material.

### 2.4. Fibrillar Collagen Quantification

Intestinal sections stained with Picrosirius red were imaged on a Zeiss Axioskop II Plus microscope with a color Zeiss Axiocam MRc camera. Three representative polarized light images per section were taken with the 10X objective using a polarizing filter. Levels were set in Zeiss Zen software and TIFF images were exported for analysis in FIJI. The sections were reviewed by an independent researcher, blinded to the treatment conditions. Images were thresholded and the percent Picrosirius red positive area was calculated. The average of all representative images per mouse was used to determine the collagen positive area/mouse. Within each experiment, imaging settings, levels, and thresholds were kept consistent to allow quantitative analysis.

### 2.5. Dermatitis and lymphedema evaluation

Dermatitis and lymphedema were assessed as previously described^23^. Details are given in Supplementary Material.

### 2.6. Microcomputed tomography (μCT) assessment of bone fractures

A 45 microCT scanner (SCANCO Medical AG, Brüttisellen, Switzerland) and 10.4 μm isotropic voxel size was used to scan blinded samples of mouse tibiae in the Department of Orthopedic Surgery of University of Pennsylvania. The resulting images were first smoothed by a Gaussian filter (sigma=1.2, support=2.0) and then a threshold corresponding to 30% of the maximum available range of image gray scale values were applied. The scanned images were 3D-reconstructed to identify the fracture sites.

### 2.7. Bioluminescence imaging of inflammation

Myeloperoxidase (MPO) activity was evaluated by bioluminescence imaging as previously described^23^. Details are given in Supplementary Material.

### 2.8. Statistical Analysis

Data are presented using means with S.E.M unless otherwise stated. Data are also presented as Box and Whisker plots showing all points (min to max), with each dot representing the average quantitative value, from a 10x or 4x field. To compare the groups, 2-sided Welch t-tests were used, and to maintain a family-wise type I error rate of 0.05, we adjusted for 3 multiple comparisons for each group of comparisons using a Holm-Bonferroni correction. To compare ordinal measurements between F-PRT and S-PRT groups (i.e., unirradiated mice had no measurable signal), the Mann-Whitney test was used. Mixed-effect models were used with a multiple comparison-Dunnett test to analyze dermatitis and chemiluminescence data from IVIS imaging, considering the interaction time and treatment between repeated measurements on the same animal. For the comparison of the Kaplan-Meier curves, Log-rank (Mantel-Cox) test was used. Statistical analysis was conducted using GraphPad Prism 9.1.2 software (San Diego, CA, USA). We performed a Fisher’s exact test that compared the frequency of fracture distribution by radiation modality and scheme.

## 3. Results

### 3.1. Reirradiation with F-PRT reduces intestinal fibrosis in a preclinical reirradiation model

We previously demonstrated that intestinal tissues are spared by F-PRT versus S-PRT as documented by the preservation of a significantly higher number of regenerated crypts and by reduction in collagen deposition and fibrosis development, in the acute and chronic phases of RT-induced damage, respectively^22, 24^. Here, we investigated whether F-PRT can similarly mitigate the long-term effects of reirradiation on whole abdomen previously irradiated with S-PRT. Mice were divided into three cohorts: i) a single dose of 12 Gy S-PRT (S-PRT); ii) a single dose of 12 Gy S-PRT followed by a single dose of 12 Gy S-PRT (S-PRT/S-PRT); and iii) a single dose of 12 Gy S-PRT followed by a single dose of 12 Gy F-PRT (S-PRT/F-PRT) (Figure 1a). At 2 months after reirradiation, all mice in irradiated cohorts developed substantial fibrosis compared to the untreated (NR) group (P<0.0001) (Figure 1b, c). The S-PRT/S-PRT-treated mice presented the highest levels of muscle layer thickening among all groups, which was significantly greater than the average muscle layer thickness in mice that received a single dose of Standard radiation (P<0.0001). In contrast, the average muscle layer thickness was significantly lower in S-PRT/F-PRT-treated mice compared to the mice from S-PRT/S-PRT cohort (P<0.0001). Interestingly, the average stromal thickness in S-PRT/F-PRT-treated mice was similar to that of mice that received a single dose of Standard radiation (S-PRT vs S-PRT/F-PRT, ns) (Figure 1b, c). Similarly, by Picrosirius red staining (Figure 1d, e), S-PRT/F-PRT treated mice had the lowest average collagen deposition after radiation, which was significantly less than the collagen deposition in S-PRT/S-PRT treated mice (S-PRT/S-PRT vs S-PRT/F-PRT, P=0.0057). Collectively, these results indicate that F-PRT ameliorates intestinal fibrosis even when applied on a previously S-PRT-treated normal intestine. As expected, there was also a decrease in the body weight of the irradiated mice following the first and the second course of PRT (Figure S1) with a weight gain that was significantly lower in both re-irradiation groups after Day 31 when compared to the single-S-PRT group (S-PRT vs S-PRT/S-PRT, P<0.0001 and S-PRT vs S-PRT/F-PRT, P<0.0001). However, the duration of periods of weight loss and the time course of recovery were very similar between the reirradiation groups (S-PRT/S-PRT vs S-PRT/F-PRT, ns).

### 3.2. F-PRT enhances survival and reduces long-term intestinal toxicity after reirradiation, including hypofractionated delivery

Next, we tested if PRT sparing of long-term RT effects on intestine can be maintained with a hypofractionated reirradiation regimen, as a more clinically relevant scenario. Mice were divided into three cohorts: i) a single dose of 12 Gy S-PRT (S-PRT); ii) a single dose of 12 Gy S-PRT followed by 3 fractions of 6.4 Gy S-PRT (S-PRT/S-PRT); and iii) a single dose of 12 Gy S-PRT followed by 3 fractions of 6.4 Gy F-PRT (S-PRT/F-PRT) (Figure 2a). Interestingly, the S-PRT/F-PRT-treated mice presented a high survival rate of 87% which was similar to the 93% survival rate of the mice treated with a single dose of S-PRT (Figure 2b). Moreover, with an 87% survival rate, the S-PRT/F-PRT-treated mice responded significantly better to reirradiation than S-PRT/S-PRT-treated mice with a 27% survival rate (P<00001) (Figure 2b), possibly due to bowel obstruction in mice reirradiated with S-PRT. To further evaluate the long-term effects of reirradiation via histological analysis, mice were euthanized at 66 days after the 3^rd^ fraction of 6.4 Gy.

Masson’s trichrome staining to assess fibrotic response revealed significant thickening of the muscle layer of all irradiated mice compared to the untreated mice (P<0.0001) (Figure 2c, d). However, the thickness of the S-PRT/F-PRT-treated intestinal stroma was significantly reduced compared to both S-PRT (S-PRT vs S-PRT/F-PRT, P<0.0193) and S-PRT/S-PRT-treated mice (S-PRT/S-PRT vs S-PRT/F-PRT, P<0.0001).

Additional assessments of collagen deposition followed a similar pattern, with collagen levels being significantly reduced after reirradiation with F-PRT compared to S-PRT (S-PRT vs S-PRT/S-PRT P<0.016; Figure e, f). Collectively, these data show F-PRT not only mitigates intestinal fibrosis relative to S-PRT, but also offers a survival benefit in a hypofractionated reirradiation scheme.

### 3.3. Reirradiation with hypofractionated F-PRT alleviates dermatitis and lymphedema

To confirm the sparing benefits of F-PRT for reirradiation of a different tissue type, we next evaluated hypofractionated reirradiation with F-PRT versus S-PRT on the skin and bone of murine leg. Radiotherapy was delivered to the leg of mice treated in three cohorts: i) a single dose of 15 Gy S-PRT (S-PRT); ii) a single dose of 15 Gy S-PRT followed by 3 fractions of 11 Gy S-PRT (S-PRT/S-PRT), given every 48hrs; and iii) a single dose of 15 Gy S-PRT followed by 3 fractions of 11 Gy F-PRT (S-PRT/F-PRT), given every 48hrs (Figure 3a). In each cohort, mice were longitudinally followed for radiation-induced dermatitis, as well as for inflammation and lymphedema over a period up to ∼3 months after the 3^rd^ fraction of reirradiation.

Radiation-induced dermatitis includes skin changes ranging from erythema and desquamation to ulceration and skin necrosis. Dermatitis was recorded three times per week based on a published system of 8 grades ranging from 0 to 3.5^25, 26^, in which a value of 1 indicates definitive erythema and a value of 3.5 indicates complete moist breakdown of the limb. Animals that received a single dose of 15 Gy (S-PRT) developed only mild dermatitis throughout the period of observation that paralleled monitoring of the reirradiation groups (i.e. for ∼3 months after reirradiation; Figure 3b). Animals that received hypofractionated irradiation following an initial single dose of 15 Gy began to develop moist desquamation on Day 10 after the second course of irradiation irrespective of whether reirradiation was with S-PRT or F-PRT. Strikingly however, treatment with S-PRT/S-PRT caused breakdown of large areas of skin with moist exudate throughout the 3-month observation period, while treatment with S-PRT/F-PRT produced only transient dermatitis. Healing took place in the 30 days after treatment with S-PRT/F-PRT, such that at 30 to 90 days after reirradiation, dermatological scores were indistinguishable from the mild dermatitis in mice that received a single dose of Standard radiation (S-PRT vs S-PRT/F-PRT, ns). In contrast, dermatitis over the 30 to 60 days after treatment with S-PRT/S-PRT was significantly worse than dermatitis after a single dose of Standard radiation (S-PRT vs S-PRT/S-PRT, P<0.0001). Kaplan-Meier plots are used to compare time to wound healing (defined as a reduction to a score of 1.25, which corresponds to dryness with some scaling upon healing) in mice reirradiated with F-PRT vs S-PRT (Figure 3c). A total of 80% of the S-PRT/F-PRT group experienced wound repair compared to only 16.6% of animals treated with S-PRT/S-PRT (S-PRT/S-PRT vs S-PRT/F-PRT, P=0.0301).

Inflammation is a common side effect of RT; thus, we monitored it after reirradiation by chemiluminescence imaging of myeloperoxidase (MPO) activity as a marker of recruited, activated myeloid cells, such as neutrophils. Myeloid cell activation peaked at 14 days after the 3^rd^ fraction of reirradiation for both the S-PRT/S-PRT and S-PRT/F-PRT groups (Figure S2a, b). This activation was higher after S-PRT/S-PRT compared to S-PRT/F-PRT, although the difference failed to reach significance. Alternatively, in a direct measure of the edema secondary to radiation induced inflammation, the width of the mouse foot was followed to monitor the onset of lymphedema, a chronic effect of RT on lymphatic vasculature leading to the accumulation of lymphatic fluid. Reirradiation with S-PRT induced lymphedema in 83.4% of mice, a significantly higher percentage than the 20% of mice that developed lymphedema after reirradiation with F-PRT (S-PRT/SPRT vs S-PRT/F-PRT, P=0.0388) (Figure 3d). Interestingly, a single course of S-PRT caused lymphedema in 12.5 % of mice, which was significantly less than the prevalence of lymphedema in mice reirradiated with S-PRT (S-PRT vs S-PRT/S-PRT, P=0.0025), but did not significantly differ from mice reirradiated with F-PRT (S-PRT vs S-PRT/F-PRT, ns) (Figure 3d).

### 3.4. Reirradiation with hypofractionated F-PRT mitigates skin fibrosis

Radiation-induced skin fibrosis is a long-term complication of RT, manifested by the deposition of fibrous connective tissue due to the accumulation of collagen and other extracellular matrix components. Here, we assessed the deposition of fibrous connective tissue in the dermal compartment of the skin by measuring its thickness in trichrome-stained positive areas (Figure 4a, b). Relative to NR mice, the thickness of fibrous connective tissue was significantly increased by a single dose of S-PRT (NR vs S-PRT, P= 0.0283) and by hypofractionated reirradiation with S-PRT (NR vs S-PRT/S-PRT, P= 0.0095). In fact, reirradiation with hypofractionated S-PRT induced the highest level of fibrotic deposition in the dermis (S-PRT/S-PRT vs S-PRT/F-PRT, P=0.0087; S-PRT/S-PRT vs S-PRT, P=0.02). In contrast, reirradiation of the skin with hypofractionated F-PRT did not significantly alter the thickness of the dermis compared to the single S-PRT group, while it closely resembled that of NR mice (S-PRT/F-PRT vs S-PRT, ns; S-PRT/F-PRT vs NR, ns).

**Figure 4.**
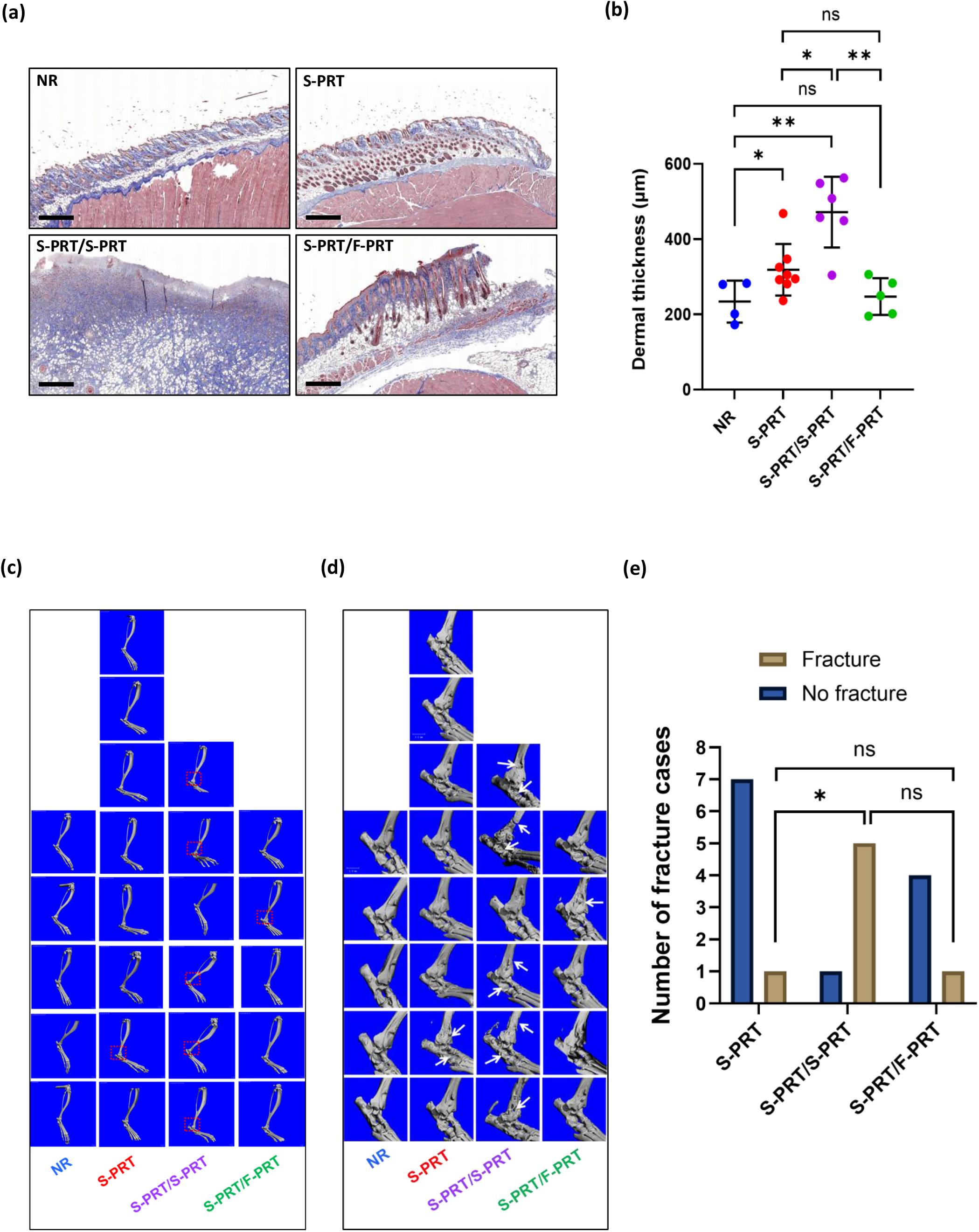
Reirradiation with F-PRT decreases the deposition of fibrous connective tissue in the skin and fracture of the tibia. **(a)** Representative images of Masson’s trichrome stain of skin sections, at 3 months after reirradiation; scale bar, 500 μm. **(b)** Scatter dot plot depicts the average dermis thickness (in μm). Each dot represents the average quantitative value per section per mouse (n=4-8). **(c)** 3D-reconstructed μCT images of mouse tibias (side view), depicting the fractured areas (red squares) at 3 months after reirradiation with the 3^rd^ fraction of 11 Gy. **(d)** Magnified areas of the μCT images (from c) presenting the damaged bone areas. **(e)** Bar graph of the number of animals with tibia fractures at 3 months after reirradiation with 3^rd^ fraction of 11 Gy. *P* values calculated with Mann-Whitney test (for b) or Fisher’s exact test (for e). In the comparison between S-PRT/S-PRT vs S-PRT/F-PRT, S-PRT/SPRT is observed as having a 13-fold increased odds of having a bone fracture (95%CI 0.60, 1160, P=0.08). *P<0.05; **P<0.01; ns, not significant. Abbreviations: NR, no-RT (control); S-PRT, Standard proton RT; F-PRT, FLASH proton RT.

### 3.5. Reirradiation with hypofractionated F-PRT protects from bone fracture

To detect fractures in the tibia of the murine leg after RT, micro-computed tomography (μCT) was employed to visualize the three-dimensional structure of leg bone. Using both a front and side view of the tibiae (Figure 4c,d and Figure S3), areas of fracture (red dotted square) were identified by a trained observer blinded to the treatment groups. In mice irradiated with a single fraction of S-PRT, 12.5% of the mice presented with fractured tibiae (Figure 4e). Reirradiation with a hypofractionated course of S-PRT produced fractures in 83.33% of mice, while reirradiation with hypofractionated F-PRT produced fracture in only 20% of the mice. In the comparison between S-PRT/S-PRT vs S-PRT/F-PRT, S-PRT/SPRT is observed as having a 13-fold increased odds of having a bone fracture (95%CI 0.60, 1160, p = 0.08). (S-PRT vs S-PRT/S-PRT, P=0.0256; S-PRT vs S-PRT/F-PRT, ns).

## 4. Discussion

To our knowledge, this is the first report on the sparing effects of FLASH RT in the context of reirradiation. Investigations are conducted on the radiosensitive normal intestine and the skin and bone of the extremities. Reirradiation with FLASH PRT after an initial scheme of Standard PRT is shown to mitigate RT-induced dermatitis, fibrosis, lymphedema, and bone damage. These acute and late toxicities are observed in the clinical setting after proton reirradiation, including in the treatment of sarcomas^27^, abdominal tumors^19, 28-30^, head and neck tumors^15, 31^, esophageal cancer^17^, breast cancer^4, 32^, and lung cancer^14, 33, 34^.

We hypothesized that F-PRT would mitigate radiation-induced fibrosis in the setting of reirradiation. It is noteworthy that our studies included evaluation of the intestine as a major dose limiting organ in RT. Patients receiving abdominal radiation can develop symptoms of radiation enteropathy, such as intestinal narrowing and transmural fibrosis after excessive submucosal accumulation of collagen and other extracellular matrix proteins, resulting in strictures and obstruction due to loss of peristaltic reflex functionality^35^. These chronic toxicities can be life altering for the long-term cancer survivor and their risk of development must be considered in planning RT for recurrent or secondary malignancies in previously irradiated abdominal areas. Surprisingly, we showed that reirradiation with F-PRT, unlike reirradiation with S-PRT, induced similar levels of collagen deposition relative to that produced by the initial dose of S-PRT. Furthermore, after reirradiation with F-PRT, intestinal muscle maintained a thickness that was indistinguishable from its thickness in the single-S-PRT irradiated group. A similar pattern was observed in fibrosis of the skin after reirradiation, thereby establishing the sparing effects of reirradiation with FLASH to include multiple tissue types.

Radiation-induced dermatitis is an additional dose-limiting toxicity of external beam RT, posing limitations in the decision-making regarding reirradiation. In the current study, reirradiation of mice with F-PRT caused dermatitis that fully resolved in 80% of mice. These data demonstrate the potential of F-PRT as a reirradiation modality that can spare radiotherapy’s late effects, such as telangiectasia and fibrosis^36^. Another late effect of radiation is lymphedema which is a chronic, inflammatory condition in which compromised lymphatic drainage leads to a local accumulation of lymph with lipids and proteins^37^. Reirradiation of the murine hind limb lymphatics with F-PRT did not significantly increase the incidence of lymphedema compared to the initial course of radiation. Crosstalk between lymphatic vasculature and skin stem cells supports wound repair^38^. Regeneration after radiation injury is also supported by lymphangiogenesis via lymphatic vessels, recently found to exist in the bone^39^. Therefore, it remains to be investigated whether the reduction in bone fractures in the F-PRT-reirradiated group relative to the S-PRT-reirradiated group relates to a persistent inflammatory microenvironment after S-PRT, as well as the possible contribution of regional lymphatic blockage, among other factors, to this response.

Although we are the first to study FLASH in the reirradiation setting, others have evaluated various approaches to delivering more than one FLASH beam in an overlapping field. Sørensen and coauthors studied how a split FLASH dose (total of 39.3 Gy) would affect murine skin toxicity on the leg. They split the dose into 2, 3, 4, or 6 deliveries with a 2-minute pause between each delivery. The data showed that the “FLASH effect” was reduced when the dose was split into 2 deliveries and abrogated when given in more than 3 deliveries^40^. In a study of RT to the lungs of nude mice bearing A549 tumors, the “FLASH effect” was maintained when radiation was delivered in 10 pulses of 2 Gy with one minute between pulses^41^. In studies of fractionated RT (without reirradiation), FLASH RT with 4 x 3.5 Gy, 2 x 7 Gy or 3 x 10 Gy delivered every 48 h of murine glioblastoma did not provoke neurocognitive decline compared to a single dose of 25 Gy^42^. Similarly, neuroprotective properties were observed by fractionated FLASH RT (2 x 10 Gy) in juvenile murine brains^43^.

Our study has several limitations. The initial dose of S-PRT in all studies was chosen to minimize acute toxicities such that reirradiation was not performed in a field already heavily damaged. If moderate toxicity did develop, mice were removed from the study. Thus, our findings are best interpreted under circumstances in which reirradiation is delivered to a field that sustained mild damage from initial RT. It should be noted that this approach is consistent with what is generally done in the clinic, where a second course of radiation would not be typically delivered to a region that currently exhibits active moderate to severe damage^44^. Nevertheless, we acknowledge that decision-making about re-irradiation is variable regarding minimum interval, contraindications and dose constraints and it is decided on case by case basis. We further note that the volume of the irradiated tissue in both intestine and skin/bone experiments is large (2 x 2 cm) and not clinically proportionate for the body size of the mice. Additional studies can be designed that will treat a smaller tissue volume. As FLASH RT has already been documented to preserve tumor response relative to Standard RT^22, 23, 45, 46^, the focus of this work is on the response of normal tissues to reirradiation with F-PRT vs S-PRT. Future studies could examine the therapeutic response of recurrent tumors to FLASH or the response of a new primary tumor that arose in a previously irradiated bed.

In conclusion, our data suggest that reirradiation of normal intestine and skin/bone tissues with F-PRT is safe and well-tolerated by mice, supporting its translational potential. Further work needs to be done to examine the radiation dose-volume relationship for the optimal use of F-PRT reirradiation and to identify the causative factors of the alleviating role of F-PRT in reirradiated epithelial and mesenchymal tissue injuries.

## Acknowledgments

The irradiations were performed by the Cell and Animal Radiation Core Facility (RRID:SCR_022377) at the University of Pennsylvania Perelman School of Medicine. We would like to thank the in-house Ion Beam Applications (IBA) physics team for facilitating the preclinical experiments. The authors acknowledge the Pathobiology Core of the Veterinary School of Medicine at the University of Pennsylvania (Pennsylvania, PA) for their invaluable contribution to this study.

