## Supplemental Material for "FLASH proton reirradiation, with or without hypofractionation, mitigates chronic toxicity in the normal murine intestine, skin, and bone"

### Supplementary information

**Murine studies**

For studies of reirradiation-induced intestinal damage, male mice (n = 8-15) were randomly assigned into three cohorts: i) a single dose of 12 Gy S-PRT (S-PRT); ii) a single dose of 12 Gy S-PRT followed in one month by a single dose of 12 Gy S-PRT (S-PRT/S-PRT); and iii) a single dose of 12 Gy S-PRT followed in one month by a single dose of 12 Gy F-PRT (S-PRT/F-PRT). In the hypofractionated reirradiation scheme of the intestine, mice (n=15) were randomly assigned to three cohorts: i) a single dose of 12 Gy S-PRT (SR); ii) a single dose of 12 Gy S-PRT followed in one month by 3 fractions of 6.4 Gy S-PRT (S-PRT/S-PRT), given every 48hrs; and iii) a single dose of 12 Gy S-PRT followed in one month by 3 fractions of 6.4 Gy F-PRT (S-PRT/F-PRT), given every 48hrs. The initial dose of 12 Gy was selected based on prior reports as a lower threshold to cause mild gastrointestinal damage[23]. For further analysis, the intestinal tissues were isolated at 2 months after reirradiation, for both irradiation schemes.

For the studies of the reirradiation-induced effects on the murine leg (hind distal femur, tibia and foot), female mice (n=4-8) were randomly divided into the following groups: i) a single dose of 15 Gy S-PRT (S-PRT); ii) a single dose of 15 Gy S-PRT followed in one month by 3 fractions of 11 Gy S-PRT (S-PRT/S-PRT), given every 48hrs; and iii) a single dose of 15 Gy S-PRT followed in one month by 3 fractions of 11 Gy F-PRT (S-PRT/F-PRT), given every 48hrs. The initial dose of 15 Gy was selected based on pilot studies in order to avoid substantial skin damage for the majority of mice in the one month period after initial RT delivery. Mice that developed moderate damage (score of 1.75) in the month after the initial dose of 15 Gy of S-PRT (5 out of 16 mice) were removed from the study. For all studies, skin and bone samples were isolated at 3 months after reirradiation.

**Proton Beam Delivery**

Briefly, proton flux was controlled by delivering a current to the beam current regulation unit. A counter was connected to an electrometer to determine pre-set counts to deliver per dose level. Absolute dosimetry was performed with a NIST-traceable calibrated Advanced Markus Chamber (PTW-Freiburg, Germany), according to the IAEA TRS-398 protocol. Online dosimetry was performed with a cross-calibrated transmission chamber (PTW Bragg Peak Chamber Type 34080, PTW-Freiburg, Germany).

**Histopathology and quantification of fibrosis**

Intestinal and skin samples were fixed in 10% formalin [Sigma, Cat#HT201128] for 24-48 hrs before dehydration and paraffin embedding. Hematoxylin and eosin (H&E), Masson's trichrome and picrosirius red stained slides were captured, using brightfield optics and a Nikon DSRi2 color camera, at 10x magnification. For each 10x magnification field, 10-15 measurements of the fibrotic stromal thickness were taken per mouse, using the Aperio ImageScope software (thickness of positive immunohistochemical staining in µm). The same procedure was followed for the quantification of skin fibrosis, acquiring the measurements from the positive trichrome-stained dermal compartment of the skin sections.

**Dermatitis and lymphedema evaluation**

Briefly, a published system of 10 grades ranging from 0.5 to 3.5 [25], was used to score the severity of dermatitis. Lymphedema was censored when the additional thickness of the irradiated leg due to swelling, caused by the accumulation of lymph after radiation, was measured to be > 1mm (using a caliper).

**Bioluminescence imaging of inflammation**

Bioluminescent images were acquired using an IVIS Spectrum imager (Perkin Elmer) from mice anesthetized with isoflurane. The images were captured 10 minutes after luminol injection when the chemiluminescence signal peaked, utilizing a 5-minute exposure time. Consistently sized regions of interest (ROI) were outlined on the irradiated right hind leg area of each mouse. The total flux (photons/sec) was quantified using Living Image Software 4.7.3 (Perkin Elmer).

**Supplementary figure captions**

**Figure S1. Weight loss of mice after intestinal reirradiation with a single dose**
**(a)** Line dot plot of the percentage weight loss of mice after single-dose of 12 Gy reirradiation of the intestine (n=8-15). *P* values calculated with 2-way ANOVA, ****P<0.0001; ns, not significant. Abbreviations: NR, no-RT (control); S-PRT, Standard proton RT; F-PRT, FLASH proton RT.

**Figure S2. Reirradiation of the hind leg with F-PRT trends with less inflammatory response. (a)** Chemiluminescence imaging of myeloperoxidase (MPO) activity is used as a marker of recruited, activated myeloid cells, such as neutrophils, in studies of inflammation. Line plots depict the total flux of chemiluminescence after reirradiation with a 3rd fraction of 11 Gy (n=3-8). **(b)** IVIS images of chemiluminescence after luminol injection. *P* values calculated with mixed-effect models, with a multiple comparison-Dunnett test. ns, not significant. Abbreviations: NR, no-RT (control); S-PRT, Standard proton RT; F-PRT, FLASH proton RT.

**Figure S3. Reirradiation with F-PRT prevents bone fractures. (a)** 3D-reconstructed µCT images of mouse tibias (front view), depicting the fractured areas (red squares) at 3 months after reirradiation with the 3$^{rd}$ fraction of 11 Gy.

### Figure S1

**(a)**

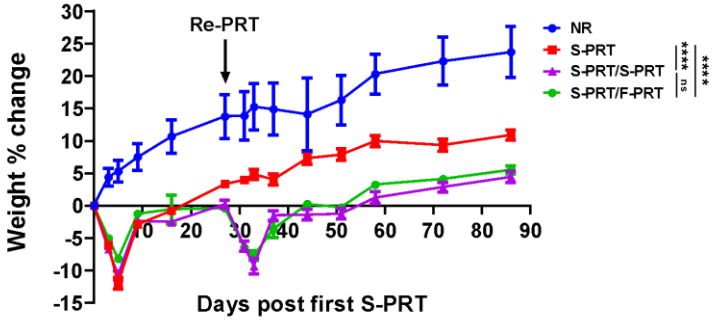
Re-PRT
Weight % change
Days post first S-PRT
NR
S-PRT
S-PRT/S-PRT
S-PRT/F-PRT
****
ns
****

### Figure S2

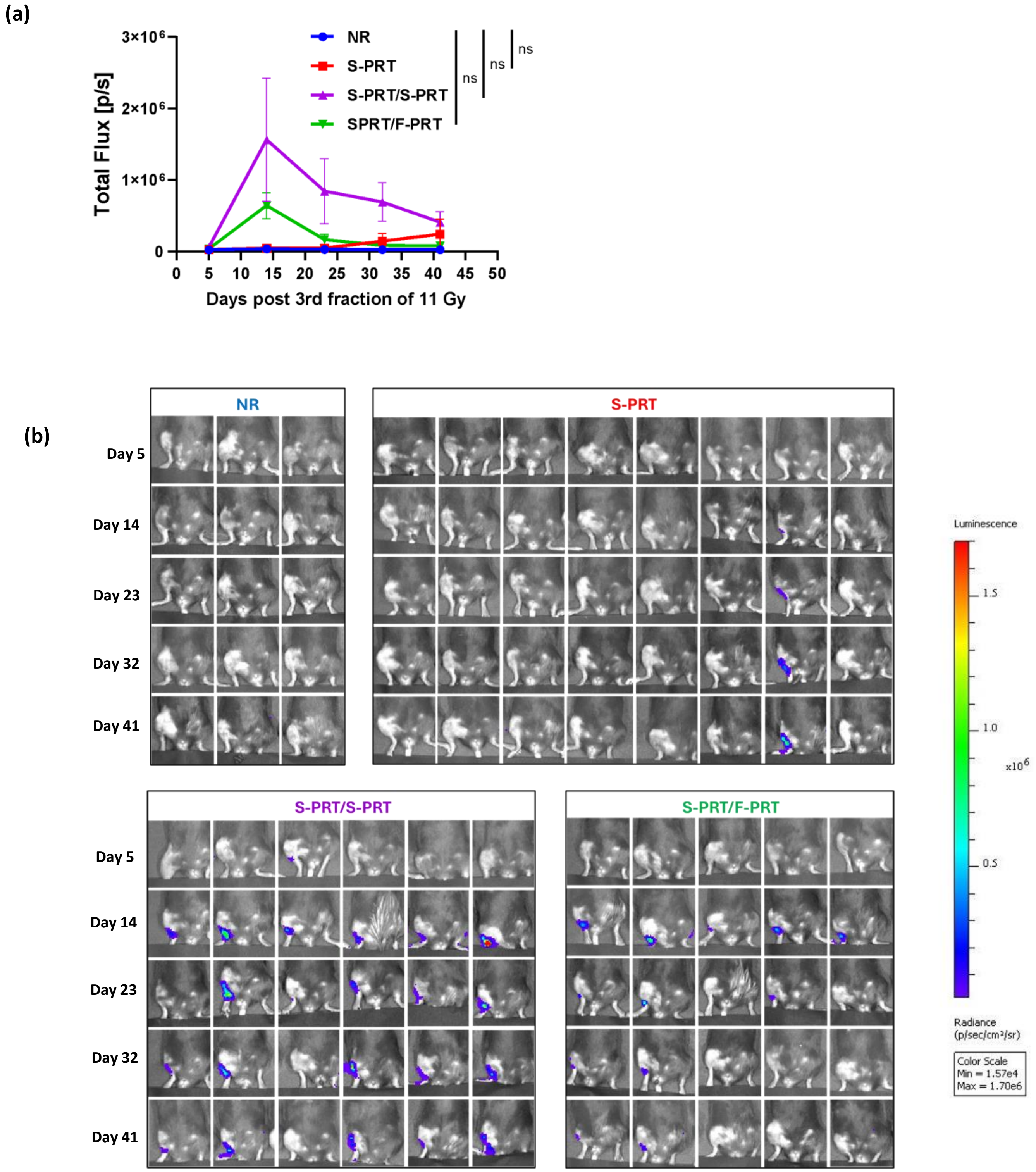
(a)
Total Flux [p/s]
3×10^6
2×10^6
1×10^6
0
NR
S-PRT
S-PRT/S-PRT
SPRT/F-PRT
ns
ns
ns
0 5 10 15 20 25 30 35 40 45 50
Days post 3rd fraction of 11 Gy
(b)
NR
S-PRT
S-PRT/S-PRT
S-PRT/F-PRT
Day 5
Day 14
Day 23
Day 32
Day 41
Luminescence
1.5
1.0
0.5
x10^6
Radiance
(p/sec/cm²/sr)
Color Scale
Min = 1.57e4
Max = 1.70e6

**Figure S3**

**(a)**

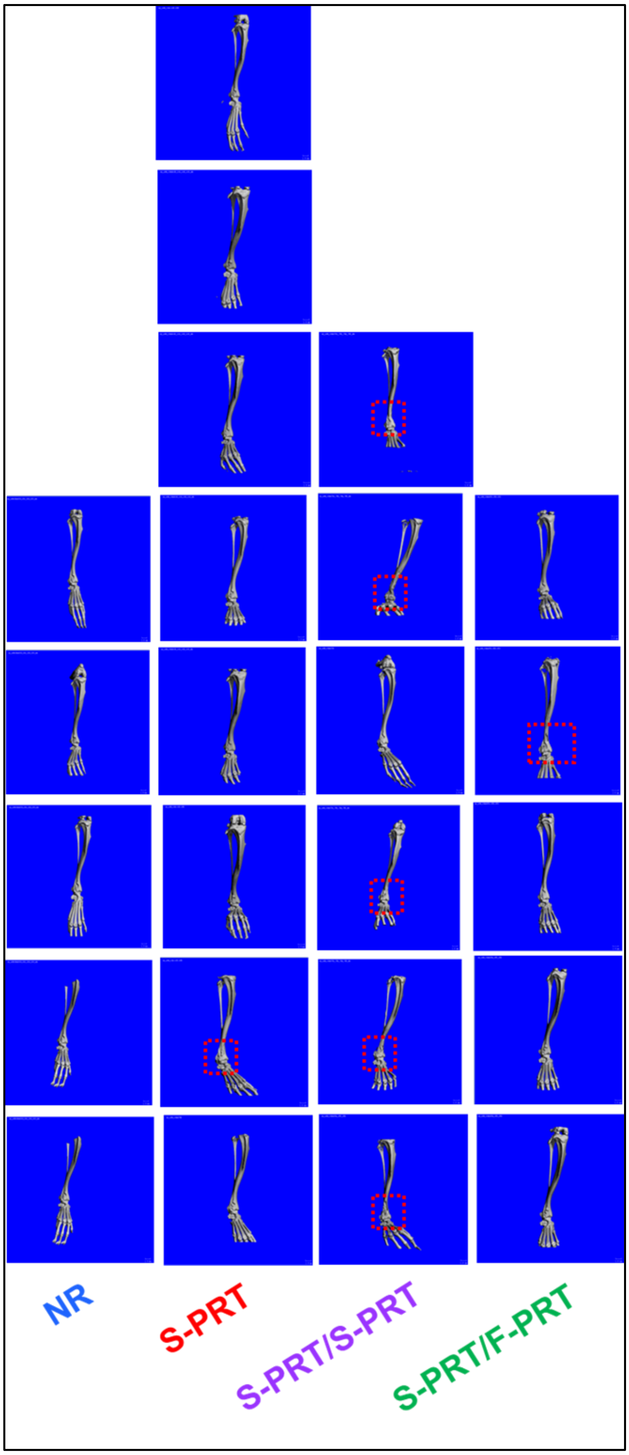
